# Deep learning approach to parameter optimization for physiological models

**DOI:** 10.1101/2025.02.25.639944

**Authors:** Xiaoyu Duan, Vipul Periwal

## Abstract

The inference of nonlinear dynamics and parameters in biological data modeling is challenging. Conventional methodologies, based on hypothetical underlying mechanisms, complicate inference because standard parameter optimization methods are difficult to constrain to biological ranges. Here, we propose a novel method to evaluate and improve putative models using neural networks to simultaneously address biological modeling, parametrization, and parameter inference. As an example, utilizing data from clinical frequently sampled intravenous glucose tolerance testing, we introduce two physiological lipolysis models (with parameters) of the dynamics of glucose, insulin, and free fatty acids (FFA). Parameter values are obtained via optimization from the limited clinical data. We then generate large quantities of simulated data from the model by sampling parameters within physiological ranges. A convolutional neural network is trained to take the simulated data time courses of glucose, insulin, and FFA as input and output the model parameters. The performance of the trained neural network is evaluated for both parameter inference and reconstruction of trajectories over a testing dataset and from optimized model-fitting curves. We show that our methodology enables accurate parameter inference and trajectory reconstruction over the testing dataset and optimized model-fitting curves. The trained neural network produces consistently high *R*^2^ values and low *p*-values across different feature engineering strategies and training dataset sizes. We assess the impact of feature engineering choices and training dataset size on inference performance, demonstrating that appropriately designed feature transformations and certain activation function improve accuracy. Our results establish a deep learning framework for parameter inference in mathematical models, which can be adapted to various physiological systems.

## Introduction

Any physiological model is based on quantitative hypotheses of a limited set of specific mechanistic interactions, with the aim of fitting model parameters to subject data in order to find subject-specific parameter values. The motivation is that deviations from specific parameter ranges may indicate a need for medical intervention, and furthermore, for clinically useful models, these may even guide the nature of a clinical intervention. Thus, estimating model parameter values from physiological data is a key problem in the development of dynamical system models of physiology.

Determining model parameters requires model optimization, especially for nonlinear models. Often, physiological parameters have specific parameter ranges, for example to ensure that predictions for plasma glucose in any model of glucose-insulin dynamics stay non-negative. Optimization algorithms are available for a large variety of problem characteristics, including constrained optimization, but here the complexity of physiology leads to issues. The underlying physiology invariably has latent processes that are not included in the model, but which lead to interactions between processes that were incorporated as independent in the model. Therefore, not just must the parameters lie within specific ranges, but they must also incorporate specific covariances that result from latent processes.

As our prior understanding of parameter space distributions given the physiological data is lacking, and the computation cost of performing forward mapping (integration) is less than that of the inverse mapping (optimization), we propose the introduction of deep learning tools to learn the inverse mapping through training neural networks. This methodology is theoretically based on the Kolmogorov universal approximation theorem, which states that in a compact region appropriate feedforward neural networks can approximate a nonlinear function [1] arbitrarily well. To facilitate neural network training, massive training data is usually required. Fortunately, in this context, generating simulated data is straightforward once the model is set up. We need to retain only parameter sets whose corresponding trajectories from integration satisfy observed physiological data constraints. While the design of the neural network can be refined through iterative trials to determine which feature engineering approach performs best, prior analysis of the underlying models can provide insights to hypothesize which features are likely to be more effective, potentially reducing the need for extensive experimentation.

In this study, we develop this deep-learning approach and apply it in detail to several ordinary differential equation (ODE) models describing insulin’s effect on lipolysis and glucose disposal as example models. The physiological models we use as examples are widely used and clinically relevant. Our aim is to provide a detailed and thorough exposition, explaining various features of our approach, that should allow others to develop such strategies for their own models.

We apply this novel approach for inferring parameter values from physiological data, utilizing frequently-sampled insulin glucose tolerance test (FSIGT) data and models of non-esterified free fatty acid (FFA) lipolysis as illustrative examples. Initially, we select several models and determine their parameter settings, followed by optimization to obtain parameter values from physiological data (including time, glucose, FFA, and insulin data), thereby establishing a distribution for each parameter corresponding to the physiological data. Subsequently, we generate sample parameter sets lying in the domains of these parameter distributions, integrating them into the model to obtain simulated data. These simulated data are partitioned into training, validation, and test datasets. The training and validation datasets are inputs to the neural network, while the corresponding parameter samples are outputs. Convolutional neural networks (CNNs) are incorporated into the architecture to capture intricate relationships between neighboring data points. During the training of such networks, we utilize the test set to evaluate the network’s performance in parameter inference, employing metrics such as reconstruction error or parameter error.

In addition to optimizing neural network architecture, our approach incorporates feature engineering to enhance the effectiveness of input data. This decision is motivated by the recognition that the relationship between raw data (such as time, glucose, FFA, and insulin levels) and outputs is likely to be more nonlinear than a neural network of a certain complexity can effectively capture in practice, as the Kolmogorov theorem applies in the limit of infinitely wide networks. As neural networks are composed of linear transformations and sigmoid functions, we introduce nonlinear transformations of the raw data as inputs to the neural network. By doing so, we aim to enable the network to capture underlying relationships without needing to be overly complex, saving considerable computational cost. The choice of nonlinear functions is informed by both empirical trials and insights derived from the ordinary differential equation (ODE) models of interest.

We demonstrate that our methodology, when applied to a well-trained neural network, achieves accurate parameter inference with high *R*^2^ values and small inference errors over the test dataset. Additionally, the reconstructed trajectories closely match the optimized model-fitting curves, confirming the reliability of inferred parameters.

Proper feature engineering improves validation loss convergence and enhances inference accuracy, particularly when combined with larger training dataset sizes. These findings highlight the effectiveness of the proposed approach in parameter estimation and model-based inference.

The paper is structured as follows: Section 1 details the physiological data and the lipolysis models used as examples. In Section 2, we outline the common procedures of our method, including the generation of simulated data, training of our proposed neural network, and parameter inference. The specifics of our approach are described in Sections 3 and 4. Section 5 presents the results, evaluating the inference performance, and the impact of feature engineering and the training dataset size. Finally, Section 6 provides a discussion of the results presented in the paper.

## 1 Data and models

### 1.1 Physiological data

In this section, we introduce two models for lipolysis mechanism to fit the physiological glucose and/or FFA data from [4], obtained through an FSIGT protocol from 25 subjects. In this protocol, glucose was administered intravenously at time zero over one minute, followed by the infusion of normal saline. Twenty minutes after the glucose injection, a bolus of insulin (0.03 U/kg) was injected. Blood samples for measuring glucose, free-fatty acid, and insulin concentrations were collected at the following minutes post-glucose injection

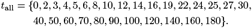

In certain models we use the time points after time 20, *t*_post20_ = {*t* ∈ *t*_all_ | *t >* 20}.We define the glucose, FFA, and insulin level measured at time *t* to be *G*_data_(*t*), *F*_data_(*t*). In particular, we denote *G*_*b*_ = *G*_data_(0), *F*_*b*_ = *F*_data_(0) and *I*_*b*_ = *I*_data_(0).

### 1.2 Two lipolysis models

In this paper, we use the following model in [3]

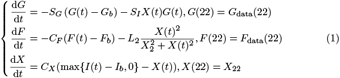

with parameter set *p* = {*S*_*I*_, *C*_*X*_, *S*_*G*_, *X*_2_, *C*_*F*_, *L*_2_}. *I*(*t*) is obtained from the insulin data *I*_data_(*t*) by linear interpolation. A remote insulin compartment *X* is introduced to account for the observed delay in insulin action on glucose. The second term in the *F* equation is a lipolysis term, which was selected in [3] from among 23 candidate models. It involves a Hill function.

[4] proposed the following novel model of FFA kinetics to study insulin action on FFA lipolysis:

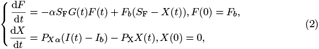

where glucose and insulin are both inputs (by linear interpolation of the discrete data over the same time course) to determine FFA clearance and lipolysis parameters, and fit to the subject data from FSIGT. They specifically designed this model to quantitatively measure the sensitivity of FFA to insulin and oxidation actions, and to estimate glucose contribution as a regulator of FFA oxidation. Here *S*_*F*_ is the fractional FFA disposal rate (1/min), *α* is a unit conversion and scaling factor for the effect of plasma glucose on the disposal of FFA, *P*_*Xα*_ is a unit conversion and a fractional transfer rate with units 1*/*min^2^, and *P*_X_ is the fractional disposal rate of insulin from the remote compartment with units (1/min). They assume that the FFA plasma concentration at time 0 equals the fasting FFA concentration *F*_*b*_.

For identifiability and to ensure that the model is linear-in-parameters, in our work we reparametrize the system (2) as

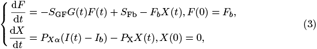

with parameter set *p* = {*S*_GF_, *S*_Fb_, *P*_*Xα*_, *P*_X_}, and will always refer to model (3) as the two-dimensional model in the remaining of this paper.

## 2 Method

Our method of inferring parameter values from physiological data requires simulated datasets for the training and testing of our neural network. As described in the introduction, each training and testing datum consists of a parameter set (the neural network output) and corresponding time-course data (the neural network input). Given physiological data from *N* subjects, traditional optimization techniques yield optimized parameter sets, resulting in *N* parameter-trajectory pairs. However, training a neural network requires a substantially larger number of such pairs. Here we introduce our methodology for generating simulated data for training purposes. It consists of the following steps, with details given in Section 3 and supplementary Text S1:

1. Optimization: Using traditional optimization techniques, estimate parameter values from the physiological data for each subject.
2. Generating non-parameter samples: Models (1) and (3) require insulin (and possibly glucose) as time-dependent inputs on the right-hand side of their X equations. To address this requirement, we employ Gaussian process regression to generate a pool of sample data for these model inputs.
3. Generating parameter samples: Use the parameter sets obtained from the optimization (Step 1) to simultaneously generate sample parameter sets and initial conditions for the ODE model.
4. Generating simulated datasets for training and testing the proposed neural network.

In Section 4, we outline the architecture and training process of our proposed neural network, referred to as the primary network to distinguish it from other smaller networks discussed in this study. The trained primary network is then used to infer parameters from denoised physiological data, followed by an evaluation of the network and model performance. We outline the process here with details in Section 4 and supplementary Text S2:

1. Feature engineering: Applied to the trajectory data to prepare the inputs for the training and testing datasets.
2. Training and testing of the network using the prepared datasets. The output of the neural network is linearly rescaled to [0, 1].
3. Evaluation of inference performance: The trained network is evaluated using two different input types: a separate testing dataset and model-fitting curves obtained from optimization. The evaluation assesses the effectiveness of the model, feature engineering, and network architecture by comparing inference performance across these two cases.

Throughout this study, we consider subjects 3, 11, 18 and 25 as outliers and exclude their physiological data from the above procedures (supplementary Text S1). The code for data generation, neural network training, and evaluation is available on GitHub at nihcompmed.

## 3 Generation of simulated data

### 3.1 Parameter optimization for models

As introduced, standard optimization methods estimate parameter values from physiological glucose and FFA data, with model integration producing system trajectories. This study aims to develop a deep learning approach to enhance the standard optimization, ensuring parameter estimations fall within physiological ranges. Optimization is still needed in the development steps, since it provides estimates of the physiological parameter ranges.

For model (1), the optimization minimizes the mean relative error between model and physiological glucose and FFA trajectories over post-20 time points. For model (3), only FFA errors are considered as the meadured glucose and insulin values are inputs to the model. Parameters are constrained to be positive by taking absolute values at each iteration, and the Nelder-Mead method is employed since it does not require the gradient. Optimization iterates until a uniform stopping criterion is met. Details and model-fitting plots are provided in supplementary Text S1.

### 3.2 Data simulation with Gaussian Process Regression

In this study, we generate simulated data (e.g., insulin for model (1)) similar to physiological data over the same time course, using Gaussian Process Regression (GPR) with a Radial Quasi-Periodic kernel. Physiological data is normalized by its mean and standard deviation at each time point, and a GPR model is trained by iteratively maximizing the marginal log likelihood between the normalized data and the model’s evaluation. At each iteration, a random subject index *i* is selected, and the model is updated until the marginal log likelihood surpasses a predefined threshold. This trained model enables the generation of simulated data over the same time course, as illustrated in Figure 1. Other details are given in supplementary Text S1.

**Fig 1.**
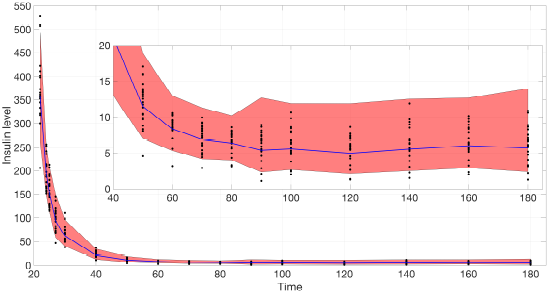
Confidence region of insulin reconstruction after *t* = 22. The blue curve is the mean predicted insulin level. The black dots are the physiological data from all 21 subjects. The inset shows a zoomed-in view of the main plot, focusing on the time range from 40 to 180 and insulin levels between 0 and 20.

### 3.3 Generation of simulated data

We use model (1) as an example to illustrate the process of generating parameter sets for simulated data. To create the simulated datasets, we require a large number of parameter sets, each corresponding to an initial condition sample set. Our goal is to use deep learning tools to infer parameter values within their physiological range. To achieve this, we impose two requirements for admissible parameter-trajectory pairs:

1. The parameters must fall within their physiological ranges.
2. The trajectories must closely resemble the shape of the model fitting curves from optimization to ensure the trained network can infer parameters accurately, performing at least as well as the optimization process.

In the remaining sections of this paper, we treat *G*_*b*_ and *F*_*b*_ as parameters of model (1), and *F*_*b*_ as a parameter of model (3). Notably, *I*_*b*_ is the first value of the insulin data generated using Gaussian process regression (GPR) as described in Section 3.2, so *I*_*b*_ does not require sampling at this stage. The initial conditions also need to be generated with the parameter samples for the integration.

We apply the following steps to generate the simulated data:

1. For each parameter, we define its physiological range to be the closed interval from the minimal to the maximal value from optimization across all subjects, excluding outliers.
2. We fit the parameter sets from optimization using a multivariate log-normal distribution over the physiological range, and generate as many parameter samples as required. This approach accounts for the correlations among different parameters.
3. We integrate each parameter set to obtain its corresponding trajectory. Notably, model (1) requires a time-dependent function *I*(*t*). For each parameter set (along with its initial conditions), we randomly select an unused insulin time-course data from the pool generated using Gaussian process regression, as described in Section For the purpose of integration, the selected insulin data is linearly interpolated over its time course.
4. We filter out the parameter-trajectory pairs whose glucose or FFA trajectory contains negative values, or do not satisfy the monotonicity condition we set. The remaining parameter-trajectory pairs serve as the training and testing data for our proposed primary neural network. The details are shown in supplementary Text S1.

Figure 2 and Figure 3 present histograms comparing the optimized parameter values (shown in red) with generated samples (shown in blue) for the 3D model (1) and the 2D model (3), respectively. In Figures 4 and 5, the simulated data are represented by blue symmetric vertical error bars, with a height equal to twice the standard deviation, for the 3D model (1) and 2D model (3), respectively. The red error bars represent the physiological data, while the black error bars represent the optimized data for both glucose and FFA in both models.

**Fig 2.**
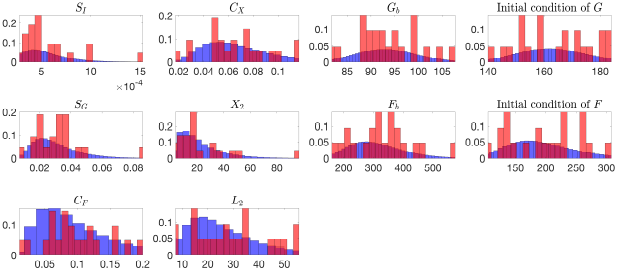
The histograms of the optimized parameter values(red) and 10,000 generated samples(blue) for the 3D model (1), both normalized to represent probabilities.

**Fig 3.**
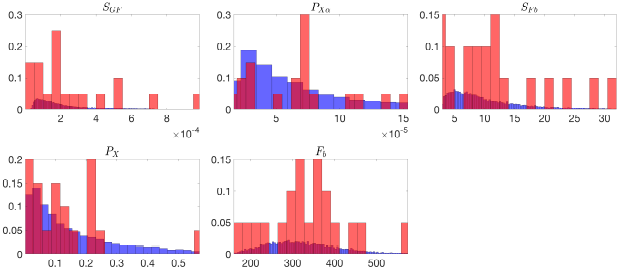
The histograms of the optimized parameter values(red) and 10,000 generated samples(blue) for the 2D model (3), both normalized to represent probabilities. Subject 4 is excluded in these histograms since its *S*_*F b*_ and *P*_*X*_ values from optimizations are much larger than the other 20 subjets.

**Fig 4.**
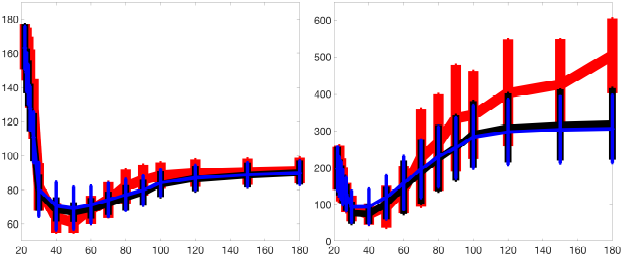
The error bars for physiological data (red), model fitting curves (black), and simulated data (blue) for glucose(left) and FFA(right) over 21 subjects with the 3D model (1).

**Fig 5.**
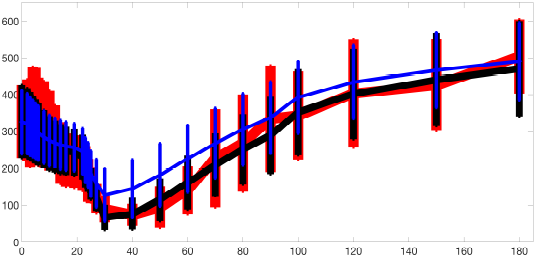
The error bars for physiological data (red), model fitting curves (black), and simulated data (blue) for FFA over 21 subjects with the 2D model (3).

## 4 Neural Network

We aim to develop a neural network that has the following inputs and outputs:

- we introduce feature engineering in Section 4.1, and apply to the simulated data we generate to obtain the neural network input *d*_*I*_, including time, glucose, FFA, and insulin information.
- the neural network output *d*_*O*_ consists of parameter sets with a size of (*N*_train_, *N*_para_), where *N*_train_ is the number of training data (in a batch or all data) and *N*_para_ is the number of parameters to be inferred by the network.

The loss function of the neural network is defined as the mean squared error (MSE) of the model evaluation based on the given outputs. A learning rate schedule, determined by the maximal learning rate, the epoch at which it is reached, and the total number of training epochs (detailed in supplementary Text S2), is used consistently for all network trainings in this study.

### 4.1 Feature Engineering and neural network architecture

The neural network we aim to train is designed to accurately capture the nonlinear relationship between the input data *d*_*I*_ (including (*t, G, I, F*) information) and the output *d*_*O*_(consisting of parameter values). According to the universal approximation theorem [1], a feedforward neural network can approximate any nonlinear function arbitrarily well within a compact region. However, in practice, the network’s ability to learn highly nonlinear relationships in raw data can be limited by computational costs and training efficiency. To overcome this challenge and improve network performance, we apply feature engineering, where we input nonlinear transformations of the (*t, G, I, F*) data into the network instead of the raw data. In this section, we describe various features of inputs, and evaluate their impact on enhancing neural network training and the accuracy of parameter inference once the training is complete.

Due to the lack of interaction between GFI(glucose, FFA and insulin) data at temporally distant time points, we put two-dimensional convolutional (conv2D) layers in our network for each feature.

To illustrate whether feature engineering has a big influence on the training and inference of the neural network, we design several features with different *d*_*I*_, but all from the same GFI data, as follows, with details shown in supplementary Text S2:

1. No Feature Engineering: *d*_*I*_ has size (*N*, 12, 16) for the 3D model (1) and size (*N*, 12, 28) for the 2D model (3). The 12 rows of *d*_*I*_ are arranged in the following order:

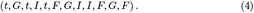

*d*_*I*_ is processed by two conv2D layers with maxpooling, and then flattened and fed into two dense layers.
2. Concatenation: the same *d*_*I*_ as in Feature 1 (No Feature Engineering) is firstly processed by a conv2D layer, whose output is concatenated with *d*_*I*_. With this feature, we put the derivatives of GFI with respect to time directly as the input of the network. The concatenated data then passes through two conv2D layers with maxpooling, is flattened, and then fed into two dense layers as before. The full process is illustrated in Figure S9.
3. Reciprocals vs. time: similar to feature 2 (Concatenation), except that *d*_*I*_ has 18 rows by attaching the following six rows to the end of (4), involving the interactions between time and reciprocals of *G, I* and *F* :

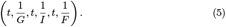
4. Mutual reciprocals: Similar to feature 2 (Concatenation), except that *d*_*I*_ has 24 rows by attaching the following 12 rows to the end of (4), involving the interactions between mutual pairs in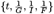:

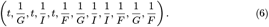

The input and output sizes and the kernel size of the conv2D layers in features 3,and 4 change correspondingly.

It is important to note that these specific sequences of reciprocals of GFI and time are not arbitrary. The point is that convolution layers have a finite kernel size and we need to make sure that all engineered features are treated equally and independently. For instance, when a conv2D kernel with size (2, 2) and stride (2, 1) moves over the first two rows of (4), it enables the neural network to approximate the time derivatives of *G* through finite difference computation. These derivatives are incorporated as part of the input to better capture the data-parameter nonlinear relationship. Reciprocals are used since *X*, the remote insulin compartment, appears in the denominator of the *F*-equation in model (1). In all features described, the activation function for all dense layers is the hyperbolic tangent function (tanh), except for the final layer, which uses either ReLU or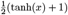 to ensure all parameters are positive before and after rescaling.

### 4.2 Training and evaluation of the inference

Now we can start to evaluate the model and the network with the introduced tools. Our primary objective is to achieve high accuracy in parameter inference from optimized model fitting curves. To this end, we independently vary the following key factors:

1. Feature engineering: for the same simulated dataset, we apply the four feature types described in Section 4.1 to obtain different inputs of the primary network.
2. Activation function: the impact of using ReLU versus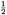(tanh(*x*) + 1) as the final activation function is assessed.
3. Dataset size: we vary the size of training and validation datasets.
4. Training hyperparameters: the effects of different batch sizes and maximal learning rates are examined.

Therefore, for any given architecture, for each training dataset size, fixing batch sizes and maximal learning rate results in eight distinct scenarios. Each scenario yields a trained network, which is subsequently evaluated based on parameter inference accuracy.

For each trained model, we apply linear regression to compute the coefficient of determination (*R*^2^) value and the *p*-value, providing a quantitative assessment of inference accuracy and statistical significance. These metrics are critical in determining the reliability of parameter estimations and guiding the selection of optimal network configurations.

## 5 Results

In this section, we evaluate the performance of the trained neural network in inferring physiological parameters. We first examine the convergence behavior of the training process and assess the inference accuracy over both the training and testing datasets, and the optimized model-fitting curves. Then, we analyze the impact of different feature engineering methods and training dataset sizes on inference performance. The results are systematically presented through validation loss curves, scatter plots of inferred versus true parameters, and comparisons of *R*^2^ values across different configurations.

### 5.1 Convergence of training and Inference Accuracy

The blue dots in Figures 6 and 7 illustrate the performance of parameter inference for eight selected parameters using scatter plots, over the training dataset (*N*_train_ = 500, 000) and testing dataset (*N*_test_ = 10, 000), respectively. Each subplot corresponds to a specific parameter (indicated in the titles), with true parameter values on the *x*-axis and inferred values on the *y*-axis. The dashed black diagonal line represents the ideal *y* = *x* relationship.

**Fig 6.**
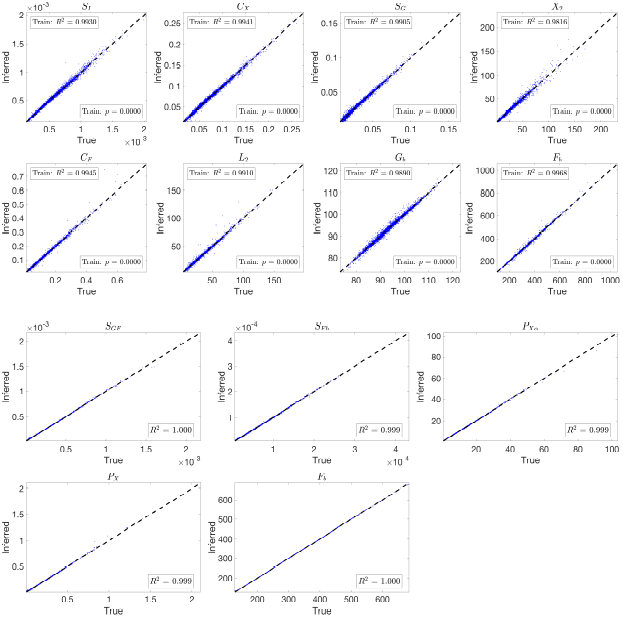
Scatter plots showing parameter inference using the trained network for the 3D and 2D models on the training dataset. Each subplot corresponds to a specific parameter, with true parameter values on the *x*-axis and inferred values on the *y*-axis. The blue dots represent the inference results, while the black dashed diagonal line indicates the ideal *y* = *x* relationship. Both networks use (tanh(*x*) + 1)*/*2 activation and the ‘Reciprocals vs. time’ feature.

**Fig 7.**
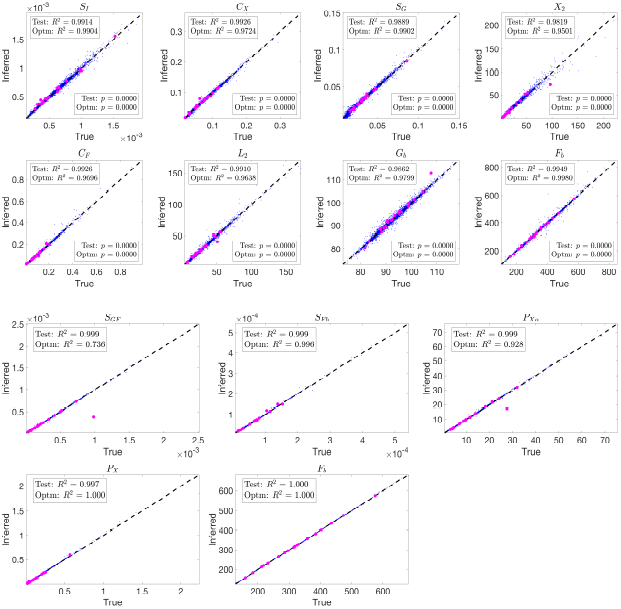
Scatter plots showing parameter inference using the trained network for the 3D and 2D models on the testing dataset and from optimized model-fitting curves. The blue dots represent inference results on the testing dataset, while the magenta dots indicate inference from optimized glucose and FFA model-fitting curves using physiological insulin data. Each subplot corresponds to a specific parameter, with true parameter values on the *x*-axis and inferred values on the *y*-axis. The black dashed diagonal line represents the ideal *y* = *x* relationship. Both networks use (tanh(*x*) + 1)*/*2 activation and the ‘Reciprocals vs. time’ feature.

To further evaluate the trained network’s ability to infer parameters from physiological data, we input optimized model-fitting curves of FFA and glucose into the trained network for each subject while using physiological insulin data directly. The optimized parameter values serve as the true values for comparison, and the resulting inference outcomes are plotted as magenta dots in Figure 7.

The inference results over both the testing dataset and the optimized model-fitting curves demonstrate high accuracy for the 3D and 2D models. The consistently high *R*^2^ values indicate a strong correlation between true and inferred parameters, while the small *p*-values confirm the statistical significance of the results. To assess whether the inferred parameters can effectively reconstruct the trajectories, we integrate the model using the inferred parameters and plot the resulting glucose and FFA curves in green in Figures S10 and S11, respectively. The close overlap between these reconstructed trajectories and the blue optimized model-fitting curves demonstrates that the trained network not only accurately infers parameters but also reproduces the model-fitting trajectories.

These results highlight the neural network’s capability to learn the relationship between input data and model parameters and to generalize effectively from simulated data to optimized model-fitting curves.

Having established the accuracy of parameter inference from both testing datasets and optimized model-fitting curves, we now investigate the factors that influence inference performance. In particular, we examine how different feature engineering strategies and varying training dataset sizes affect both validation loss and *R*^2^ values. This analysis provides insights into how architectural choices impact the model’s learning efficiency and inference accuracy.

### 5.2 Effect of Feature Engineering and Impact of Training Data Size

To further quantify the effect of feature engineering and training dataset size, we compute the *R*^2^ values across all inferred parameters. By comparing these values across different conditions, we can systematically assess how inference accuracy evolves as training progresses and whether incorporating additional features leads to more robust predictions.

Figure 8 and 9 present the validation loss curves over 2000 epochs for different feature engineering methods and training dataset sizes, for the 3D and 2D models respectively. Each subplot corresponds to a specific training dataset size (*N*_train_), increasing from left to right, with the top row using the ReLU activation function and the bottom row using the (tanh(*x*) + 1)*/*2 activation function. The four curves in each subplot represent different feature engineering strategies: No Feature Engineering (red), Concatenation (green), Reciprocals vs. Time (blue), and Mutual Reciprocals (black). Insets zoom in on the final 500 epochs to highlight convergence trends.

**Fig 8.**
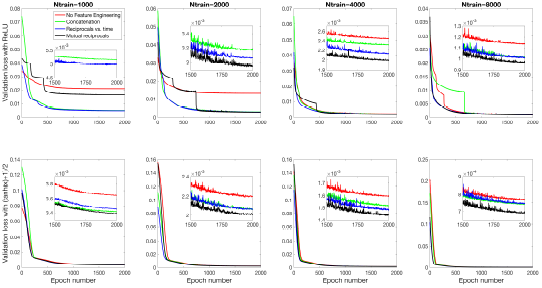
Validation loss curves over 2000 epochs for different feature engineering methods and training dataset sizes for the 3D model. Each subplot corresponds to a specific training dataset size (*N*_train_), increasing from left to right, with the top row using the ReLU activation function and the bottom row using the (tanh(*x*) + 1)*/*2 activation function. The four curves in each subplot represent the different feature engineering strategies: No Feature Engineering (red), Concatenation (green), Reciprocals vs. Time (blue), and Mutual Reciprocals (black). Insets zoom in on the final 500 epochs to highlight convergence trends.

**Fig 9.**
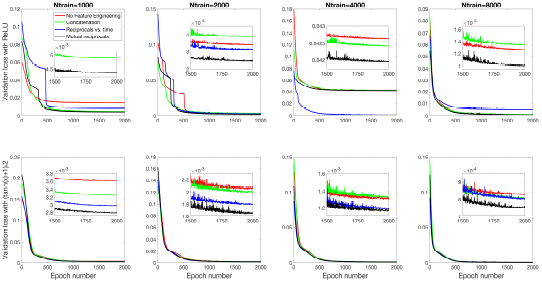
Validation loss curves over 2000 epochs for different feature engineering methods and training dataset sizes for the 2D model. The notations are similar to Figure 8.

Figures 10 and 11 present the mean *R*^2^ values as a function of training dataset size for the 3D and 2D models, respectively, under different feature engineering methods and activation functions. The upper two subplots in each figure show results over the same fixed testing dataset, while the lower two subplots show results from optimized model-fitting curves. The consistency of the testing dataset across models allows for a fair comparison of effect of feature engineering and training dataset sizes.

**Fig 10.**
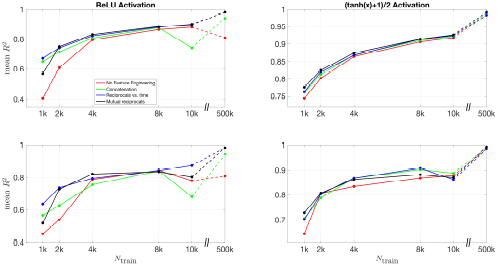
Mean *R*^2^ values for different feature engineering methods as a function of training dataset size in the 3D model. The upper two subplots show results over the same fixed testing dataset, while the lower two subplots show results from the optimized model-fitting curves. The left column corresponds to ReLU activation, while the right column corresponds to (tanh(*x*) + 1)*/*2 activation. Each color represents a different feature engineering method: No Feature Engineering (red), Concatenation (green), Reciprocals vs. Time (blue), and Mutual Reciprocals (black). The dashed lines connect the results at *N*_train_ = 10, 000 and *N*_train_ = 500, 000, illustrating the increasing trend in *R*^2^ values as training dataset size grows.

**Fig 11.**
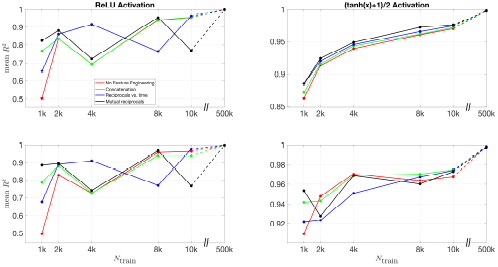
Mean *R*^2^ values for different feature engineering methods as a function of training dataset size in the 2D model. The upper two subplots show results over the same fixed testing dataset, while the lower two subplots show results from the optimized model-fitting curves. The left column corresponds to ReLU activation, while the right column corresponds to (tanh(*x*) + 1)*/*2 activation. Each color represents a different feature engineering method: No Feature Engineering (red), Concatenation (green), Reciprocals vs. Time (blue), and Mutual Reciprocals (black). The dashed lines illustrate the improvement from *N*_train_ = 10, 000 to *N*_train_ = 500, 000, demonstrating that as the training dataset size increases, the mean *R*^2^ values also improve across all feature engineering methods.

For *R*^2^ values computed for each parameter across different features and training dataset sizes, Figures S12 and S13 present results for the 3D model over the testing dataset and optimized model-fitting curves, respectively, while Figures S14 and S15 provide the corresponding results for the 2D model. These figures offer a detailed view of how each individual parameter’s inference accuracy varies with training data size and feature engineering choices. All four figures use the (tanh(*x*) + 1)*/*2 activation function.

For the (tanh(*x*) + 1)*/*2 activation function, the improvement in *R*^2^ values is more pronounced as more features are included. Across all dataset sizes, models using feature engineering techniques such as concatenation, reciprocals vs. time, and mutual reciprocals generally perform better than models without feature engineering. However, for smaller training dataset sizes, there are some fluctuations in performance, particularly for ReLU activation. Despite these variations, with a sufficiently large training dataset, the *R*^2^ value consistently improves across almost all feature engineering methods, highlighting the significant impact of dataset size on inference accuracy. The dashed lines illustrate this substantial improvement as training dataset size increases.

### 5.3 Conclusion

The results presented in this section demonstrate the effectiveness of the trained neural network in accurately inferring physiological parameters. The evaluation over both the training and testing datasets, as well as the optimized model-fitting curves, confirms that the network consistently achieves high *R*^2^ values and statistically significant *p*-values, indicating strong inference accuracy.

Further analysis of feature engineering methods and training dataset sizes highlights key factors influencing model performance. The validation loss curves reveal that feature engineering generally improves training efficiency, particularly when using the (tanh(*x*) + 1)*/*2 activation function. The comparison of *R*^2^ values across different configurations shows that models trained with feature engineering methods, such as mutual reciprocals and reciprocals vs. time, tend to achieve better inference accuracy than models without feature engineering. While performance fluctuations are observed for smaller training datasets, the overall trend indicates that increasing the dataset size leads to improved inference accuracy across all feature engineering methods. Notably, with a sufficiently large training dataset, the *R*^2^ values consistently reach higher levels, demonstrating the importance of dataset size in achieving robust parameter inference.

Additionally, inference results from the optimized model-fitting curves further validate the network’s ability to generalize beyond simulated data. The close match between the reconstructed trajectories and the original optimized model-fitting curves confirms that the inferred parameters are not only accurate but also capable of reproducing the expected physiological dynamics.

These findings highlight the strength of the proposed neural network approach in parameter inference, demonstrating its potential for broader applications in modeling complex physiological systems. Extending this framework to explore other physiological models will require custom feature engineering strategies based on the specific characteristics of the model, enabling more tailored and effective parameter inference.

Our method also has potential for early detection of potential parameter identifiability issues. In cases where multiple parameter sets generate similar trajectory outputs, the trained network may identify alternative parameter sets from the data.

Even if the inferred parameters deviate from expected values, the reconstructed trajectories remain close to the target. This property suggests that the neural network could serve as a tool for detecting and analyzing non-identifiability issues in complex physiological models. Identifying such cases is crucial for refining model structures and improving their predictive power.

While these results establish a strong foundation for parameter inference, an important next step remains: inferring parameters directly from (denoised) physiological data instead of optimized model-fitting curves. These two data sources are not identical, as model-fitting curves are generated based on mathematical models that, while descriptive, may not fully capture the complexity of real physiological systems. Limitations in the model can introduce discrepancies between the optimized curves and physiological data over certain time courses. A key challenge is to develop a mapping that transforms real physiological data into curves that retain the essential characteristics of model-fitting curves, allowing the trained network to infer parameters with similar accuracy. Addressing this challenge will require additional efforts in model refinement, feature engineering, and data processing to bridge the gap between real-world measurements and model-based inference.

In supplementary Text S2, we propose a denoising method by training a separate neural network that learns to transform raw physiological data into smoother curves that closely resemble model-fitting curves. This denoising model is trained on simulated noisy data paired with corresponding model-fitting curves, enabling it to remove noise and retain the underlying dynamics. By preprocessing physiological data using this approach, we aim to reduce the discrepancy between real-world data and model-generated curves, improving the robustness of parameter inference from physiological measurements.

## 6 Discussion

### 6.1 Limitation of current approach

1. While the models in this study show stable convergence and accurate inference, applying the framework to more complex physiological systems introduces additional computational challenges. As model complexity increases, achieving low validation loss and accurate inference for all parameters requires more computing resources and longer training times. More complex models may also need extra techniques to prevent overfitting to the simulated training data. Ensuring reliable performance in such systems remains an area for further study.
2. One challenge in using this framework with real-world physiological data is converting noisy experimental measurements into model-compatible inputs. Although supplementary Text S2 introduces a denoising method, further improvements are needed to ensure the denoised data closely match model-generated trajectories. Developing a reliable way to transform real data into model-like inputs is important because differences between them can lead to inference errors. Addressing this issue is key to applying this method beyond simulations to real-world clinical and physiological data.
3. The insulin values are currently sampled independently from the pool generated by Gaussian Process Regression (GPR) during data generation. While this approach has not caused significant issues in our current results, it may introduce inconsistencies when capturing dependencies between insulin and other physiological variables. Addressing this limitation in future work could further refine the accuracy and robustness of the inference process.

## Acknowledgments

This research was supported [in part] by the Intramural Research Program of the NIH, The National Institute of Diabetes and Digestive and Kidney Diseases (NIDDK), grant ZIA Z01-DK013030.

---

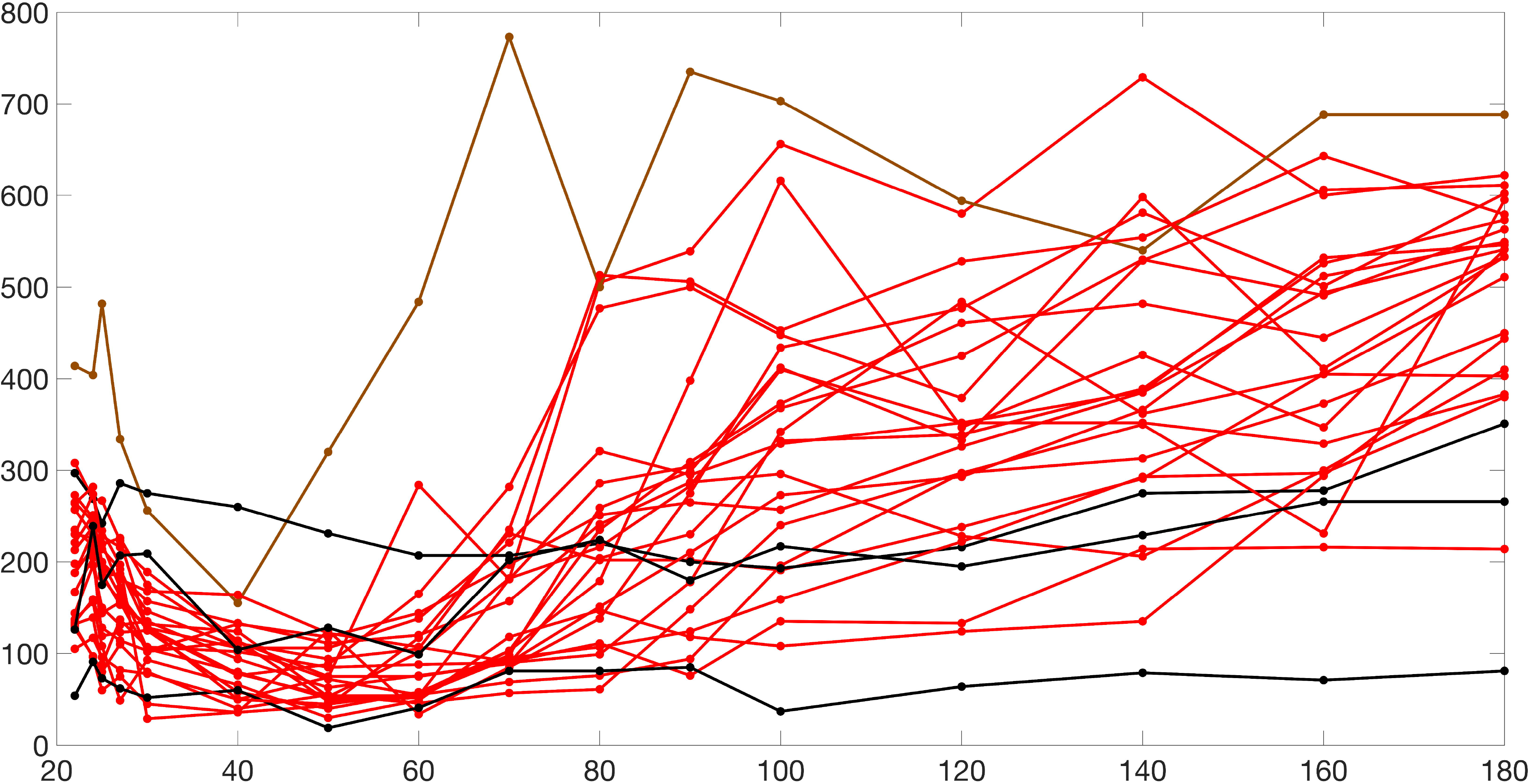

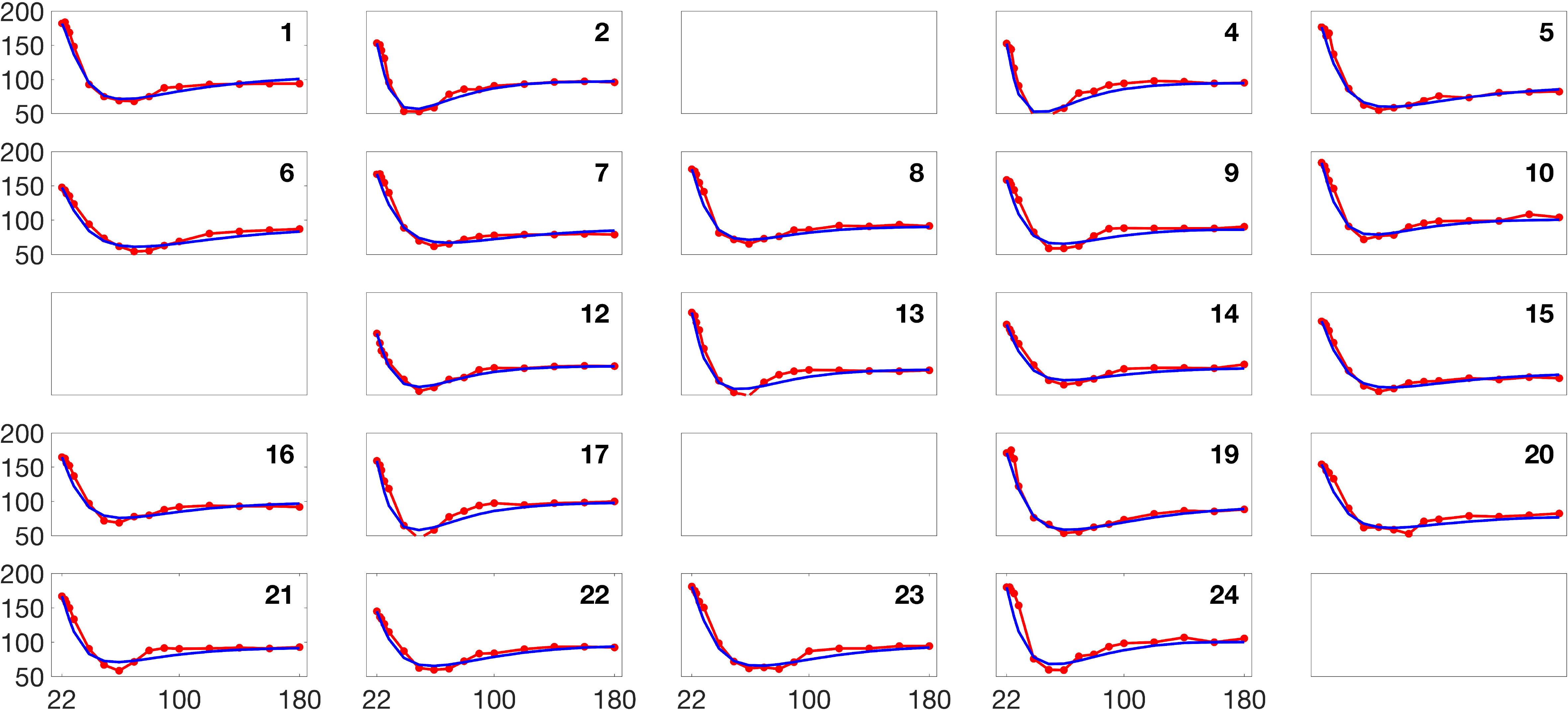

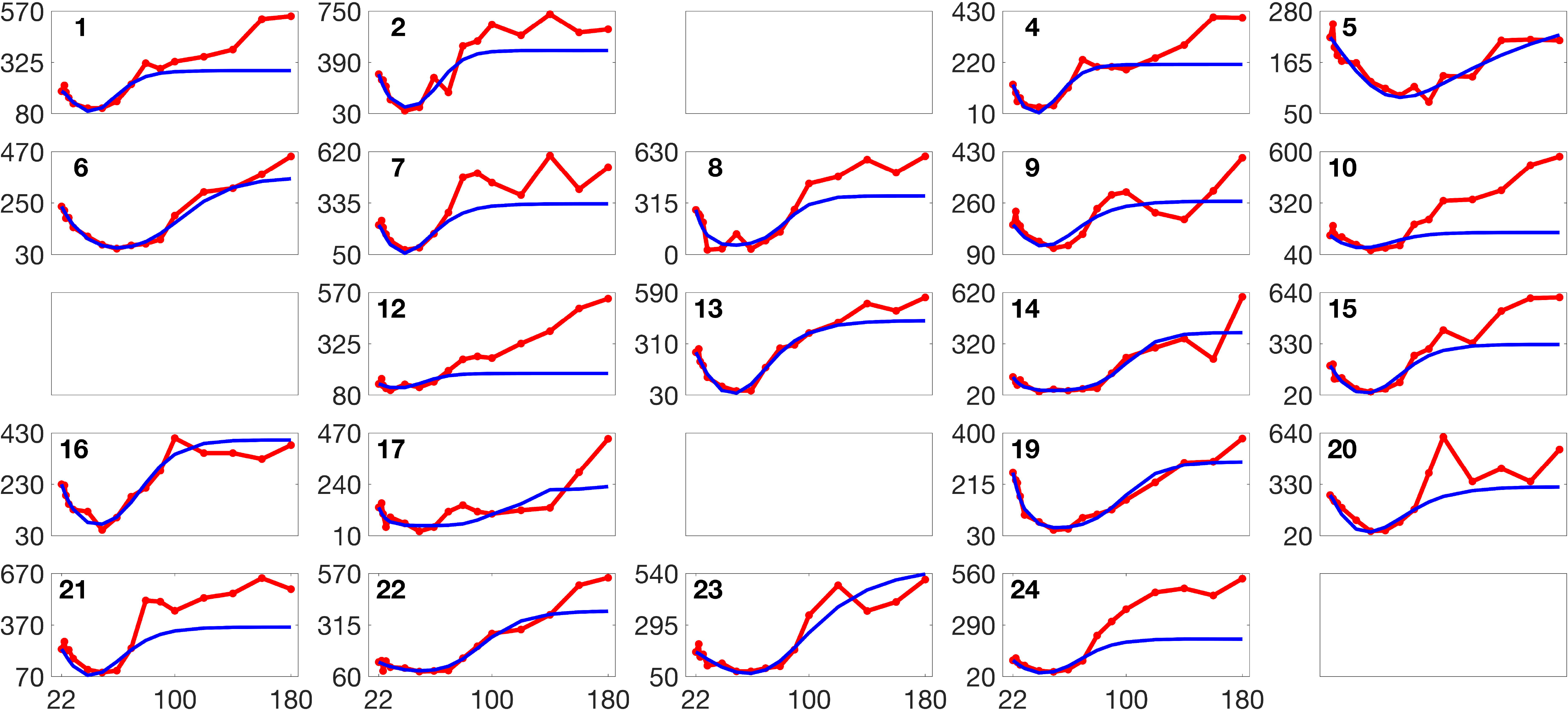

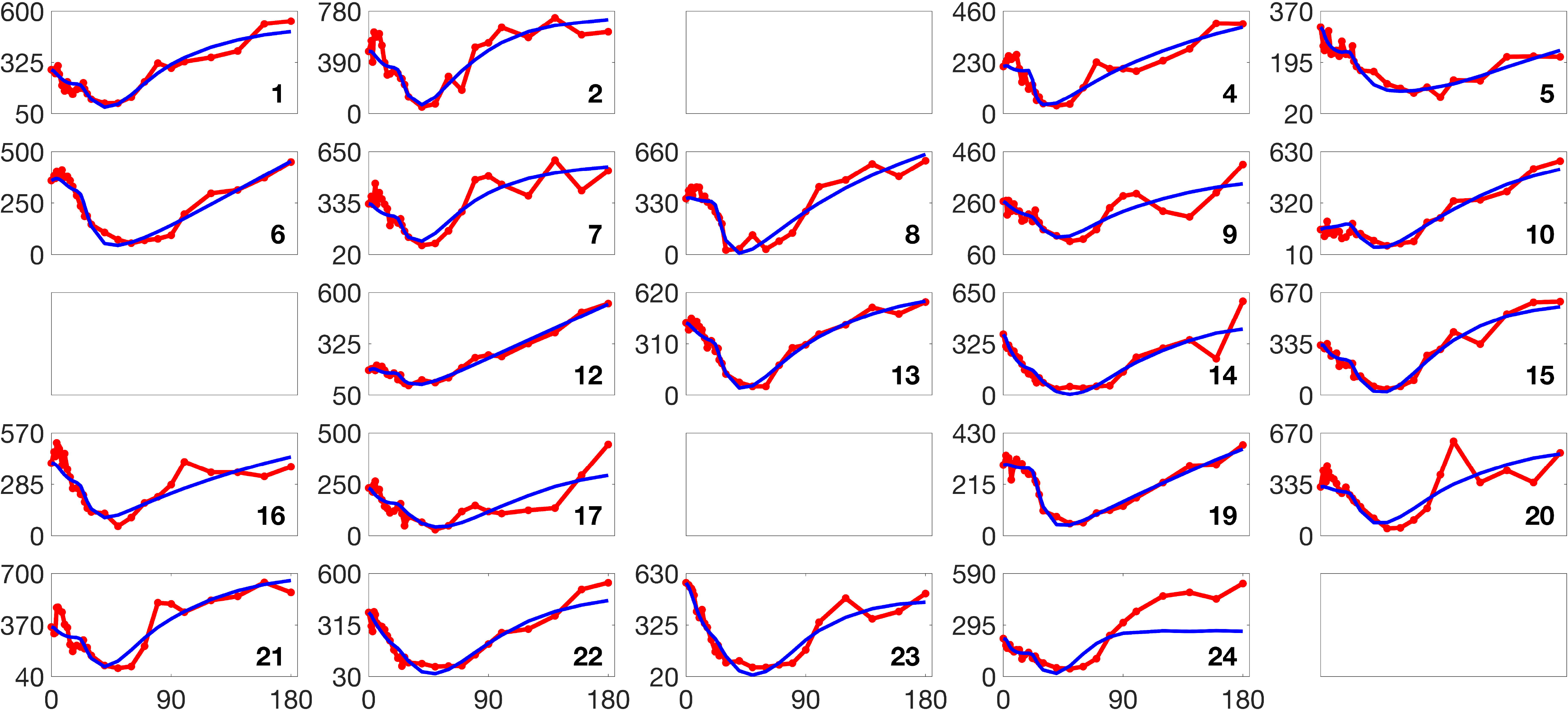

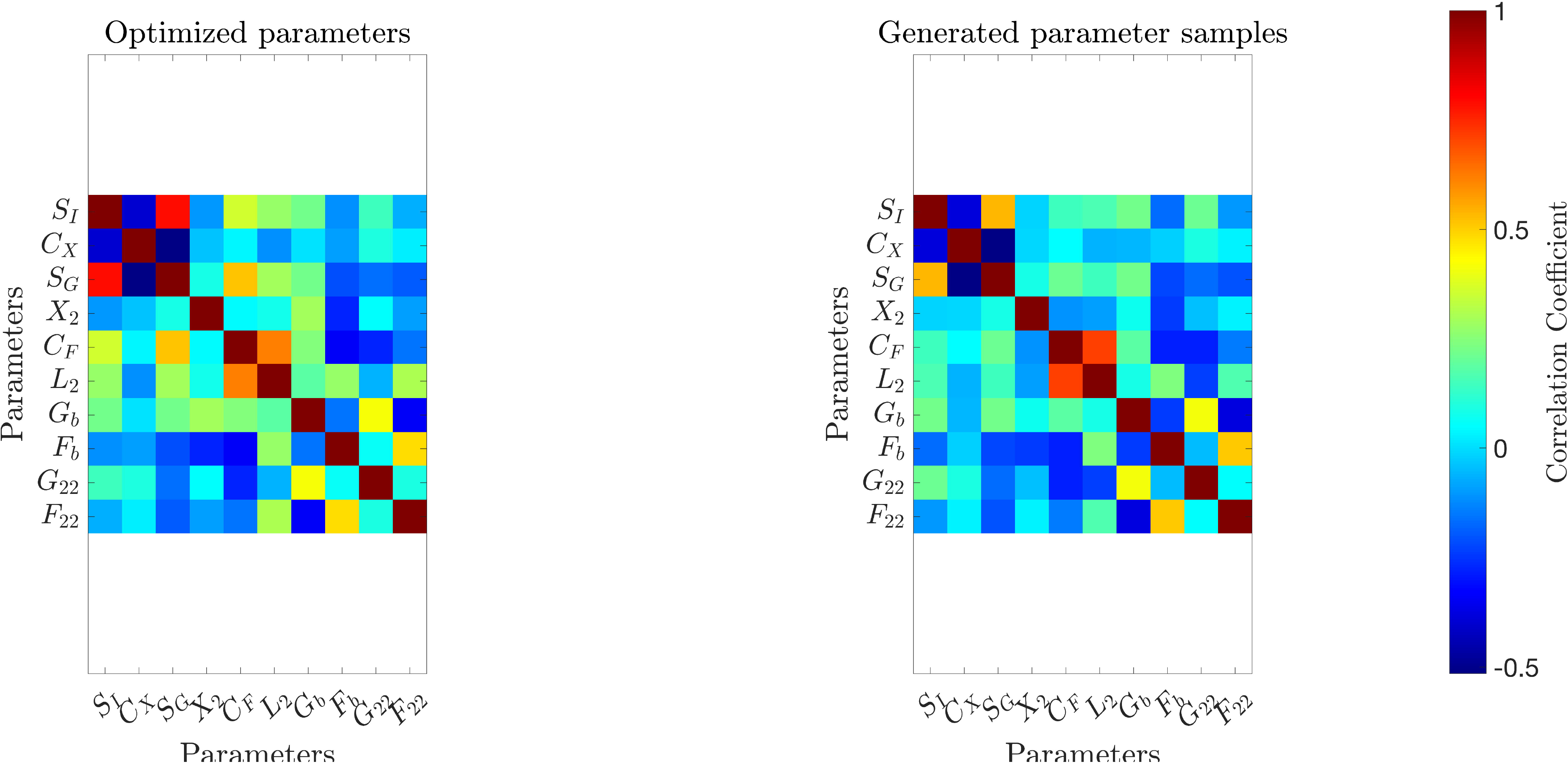

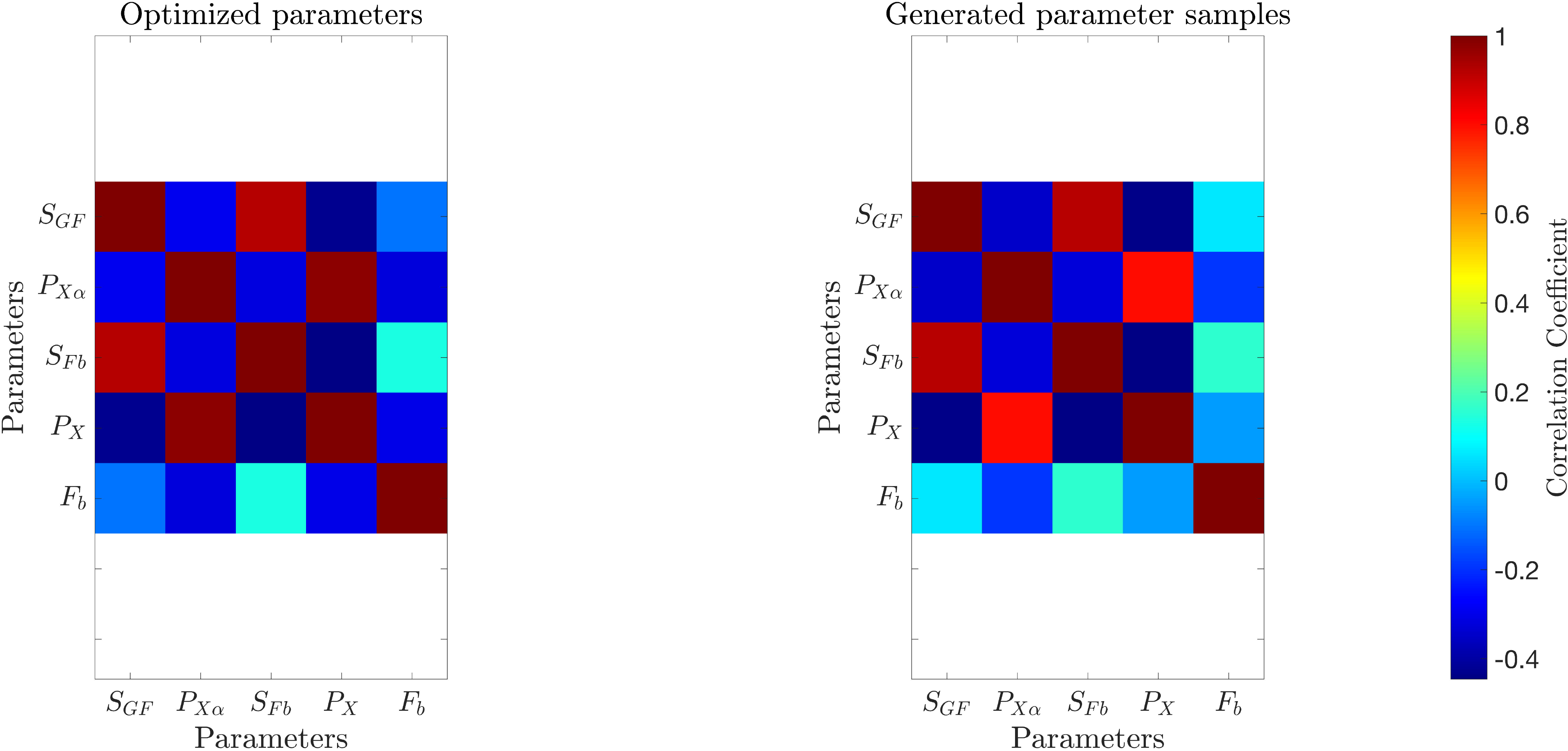

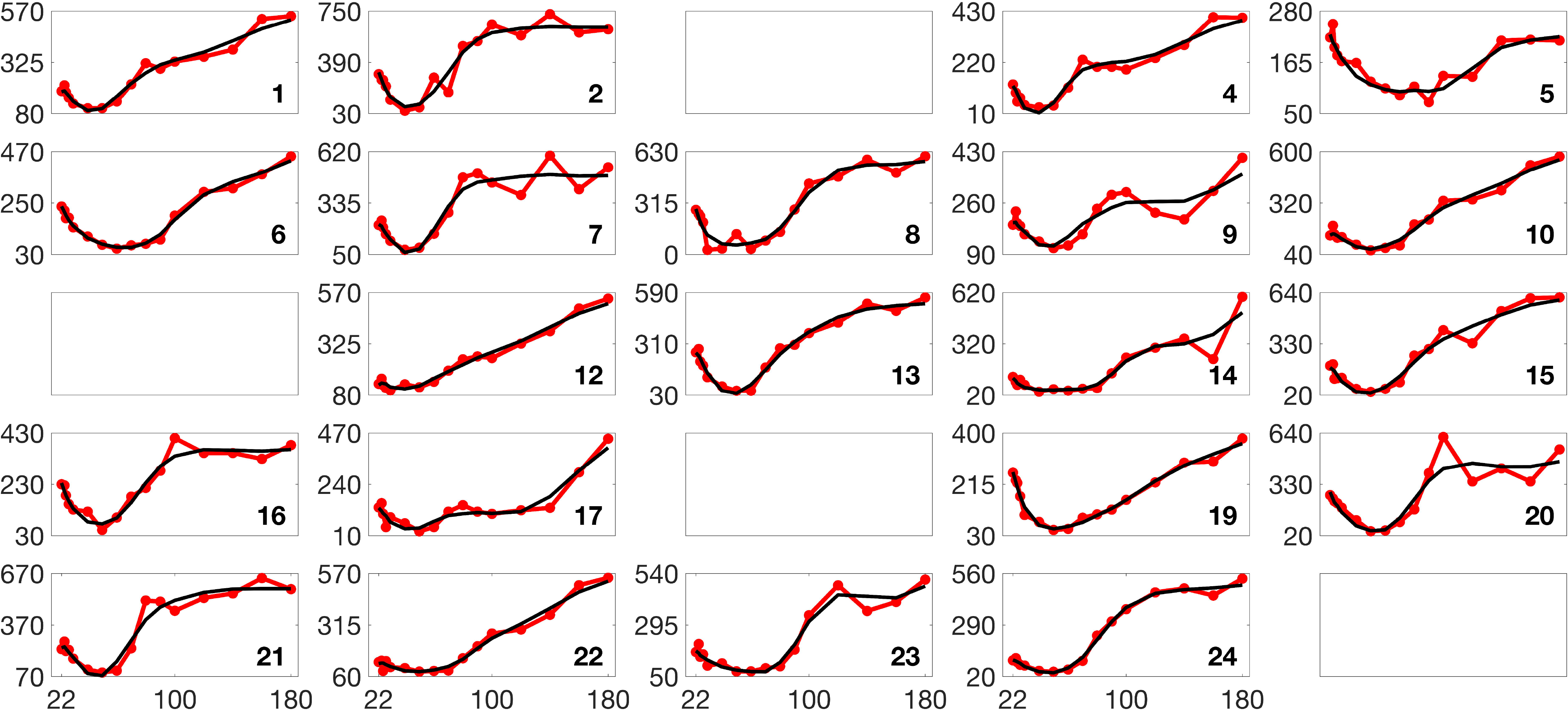

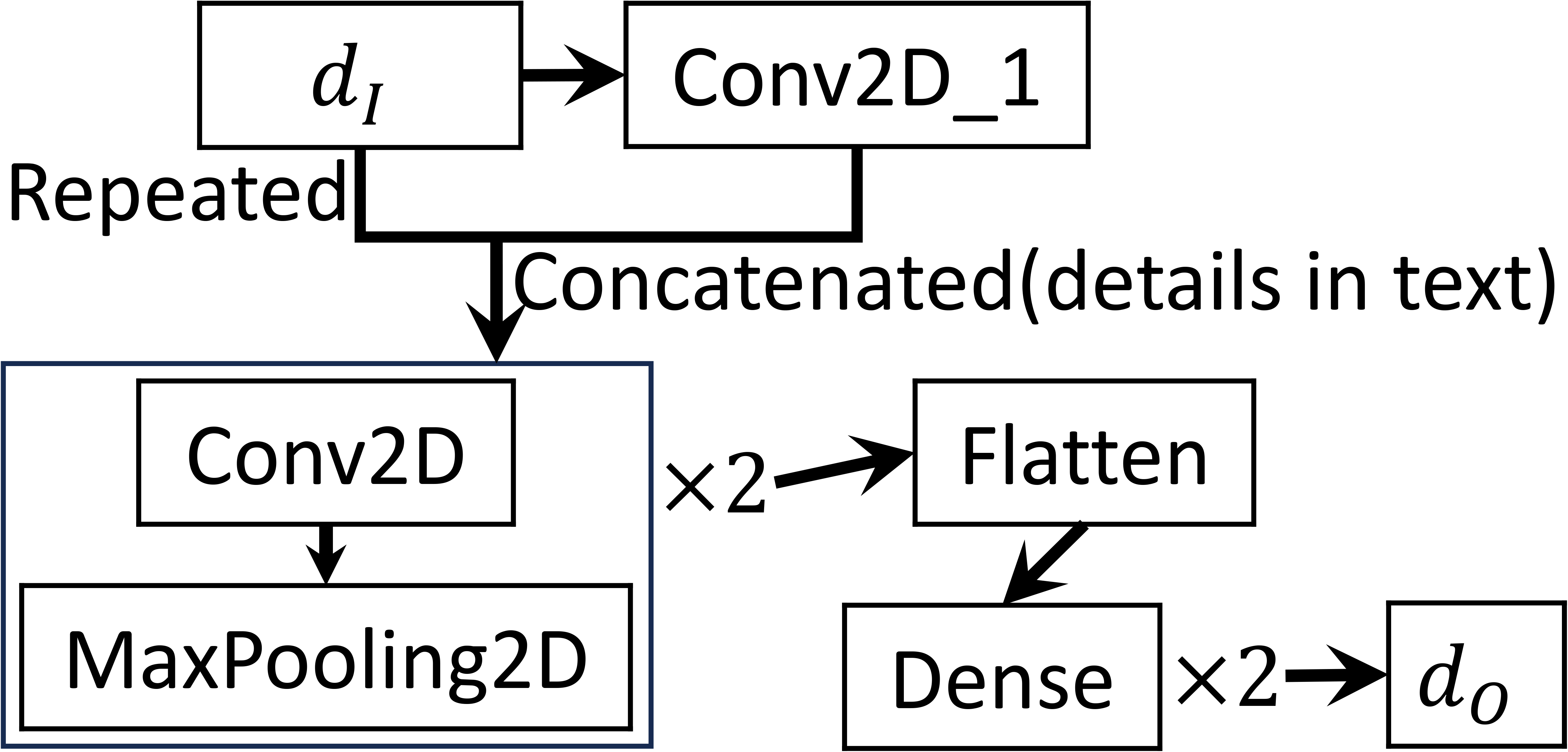

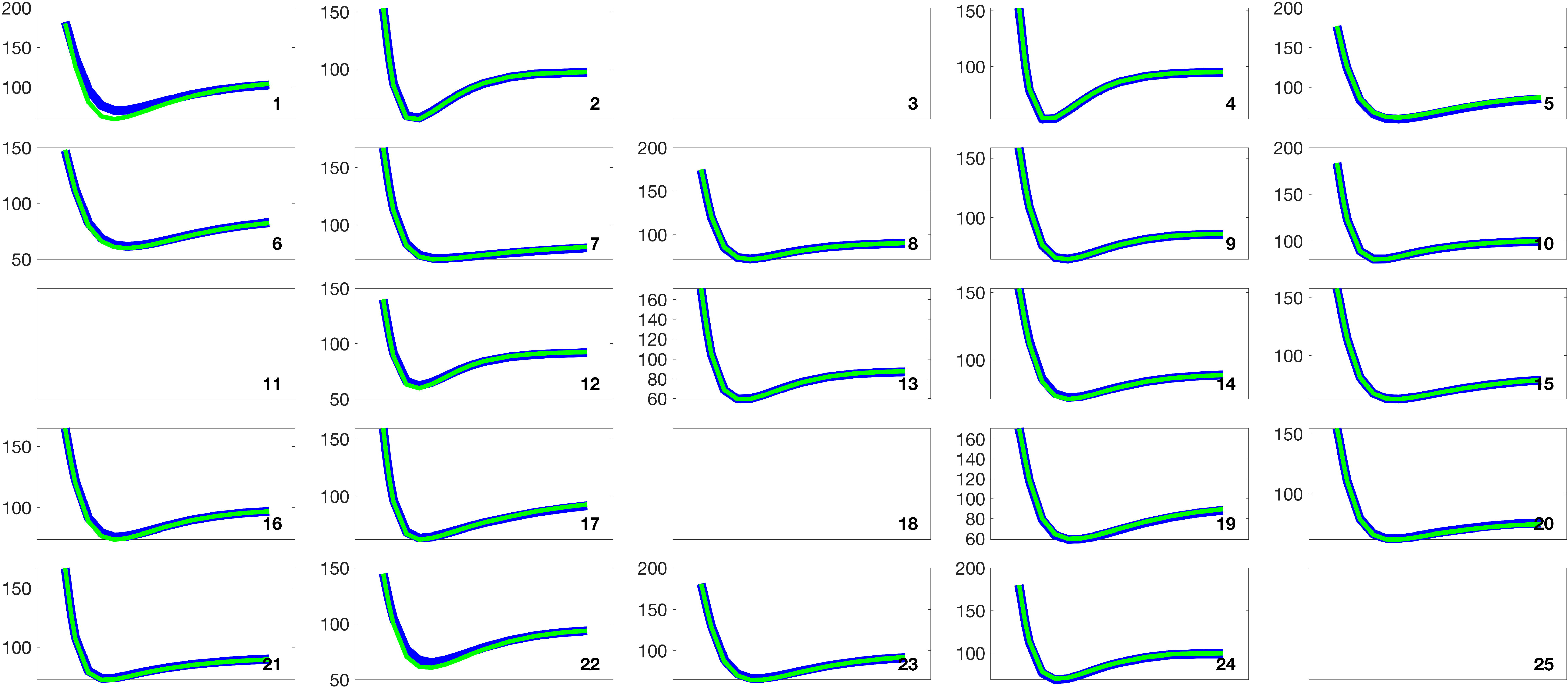

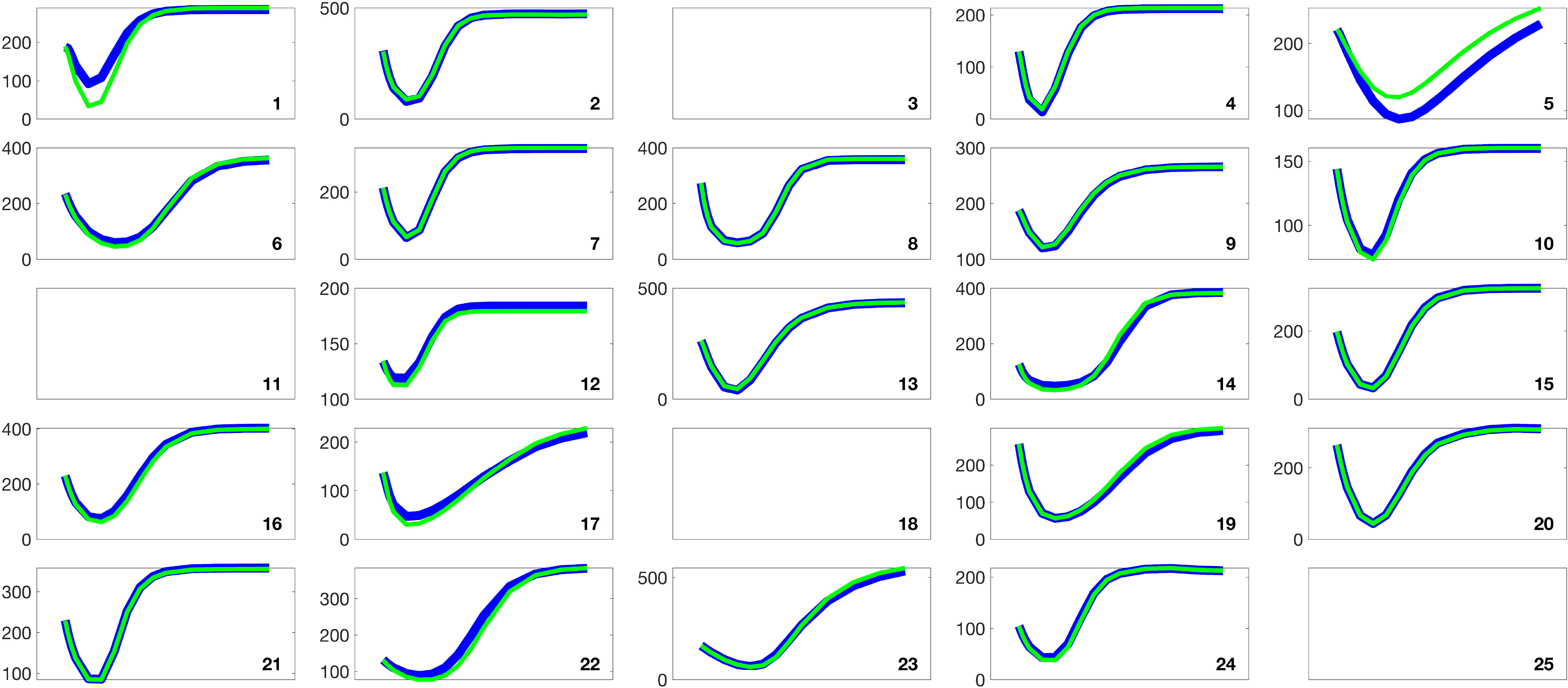

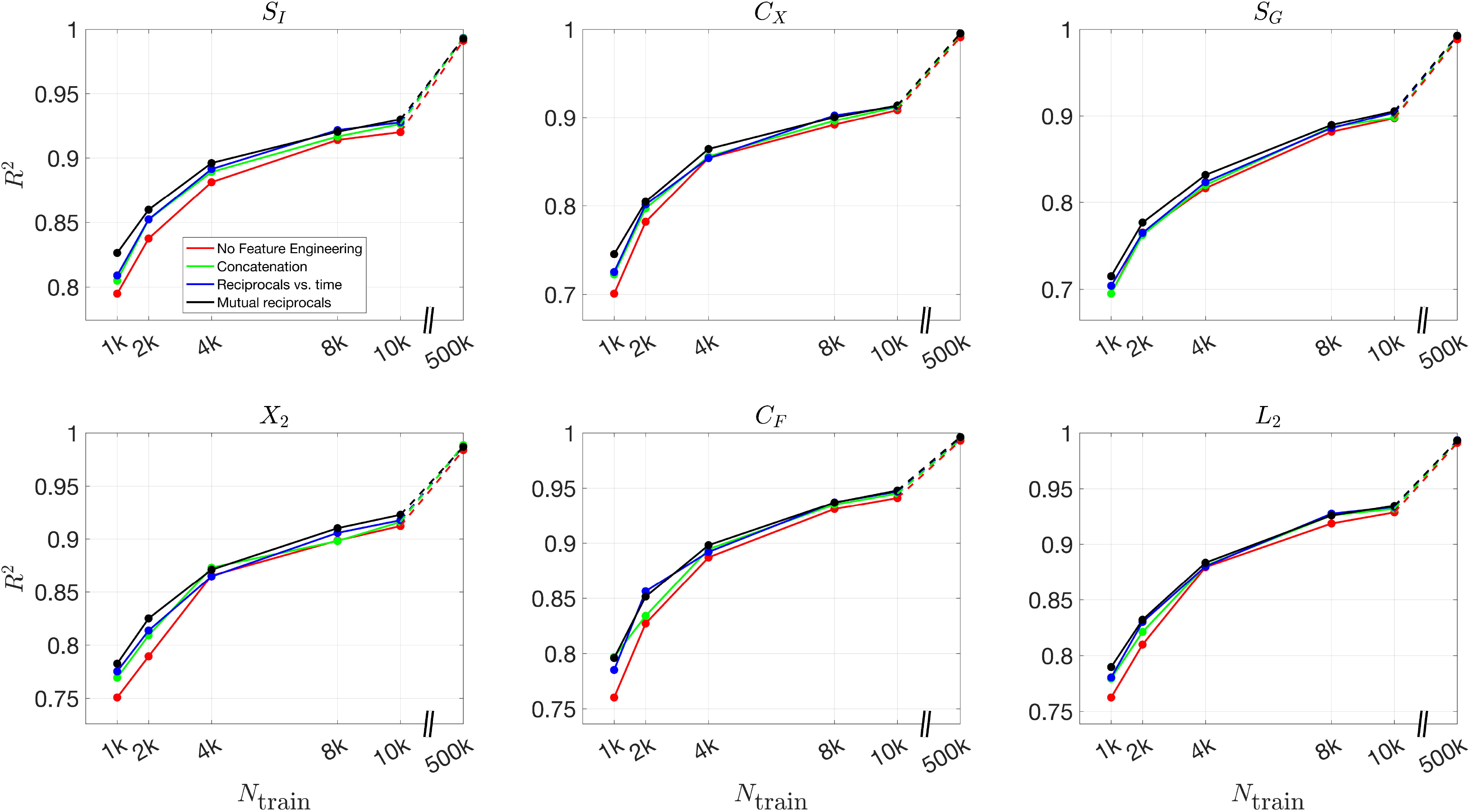

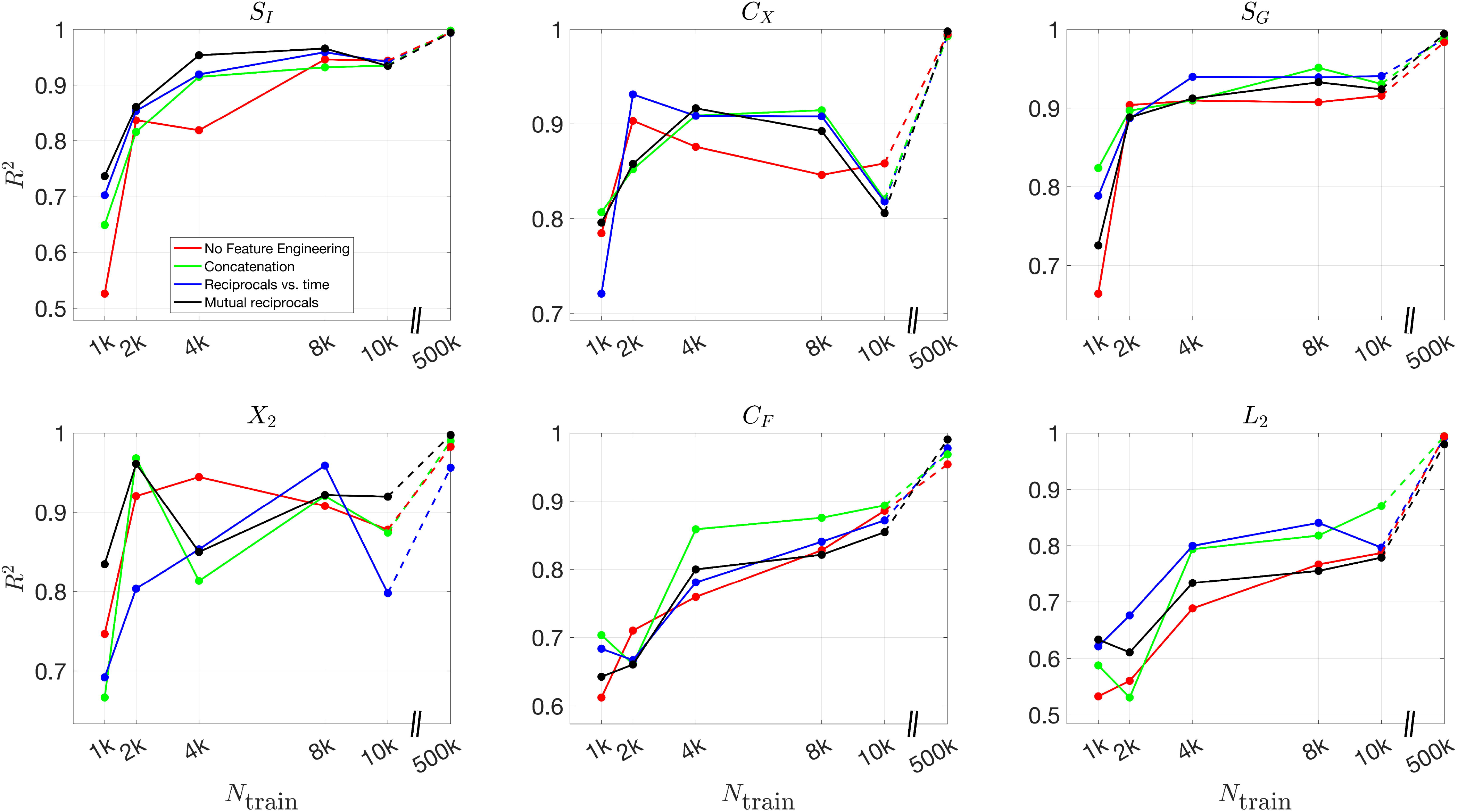

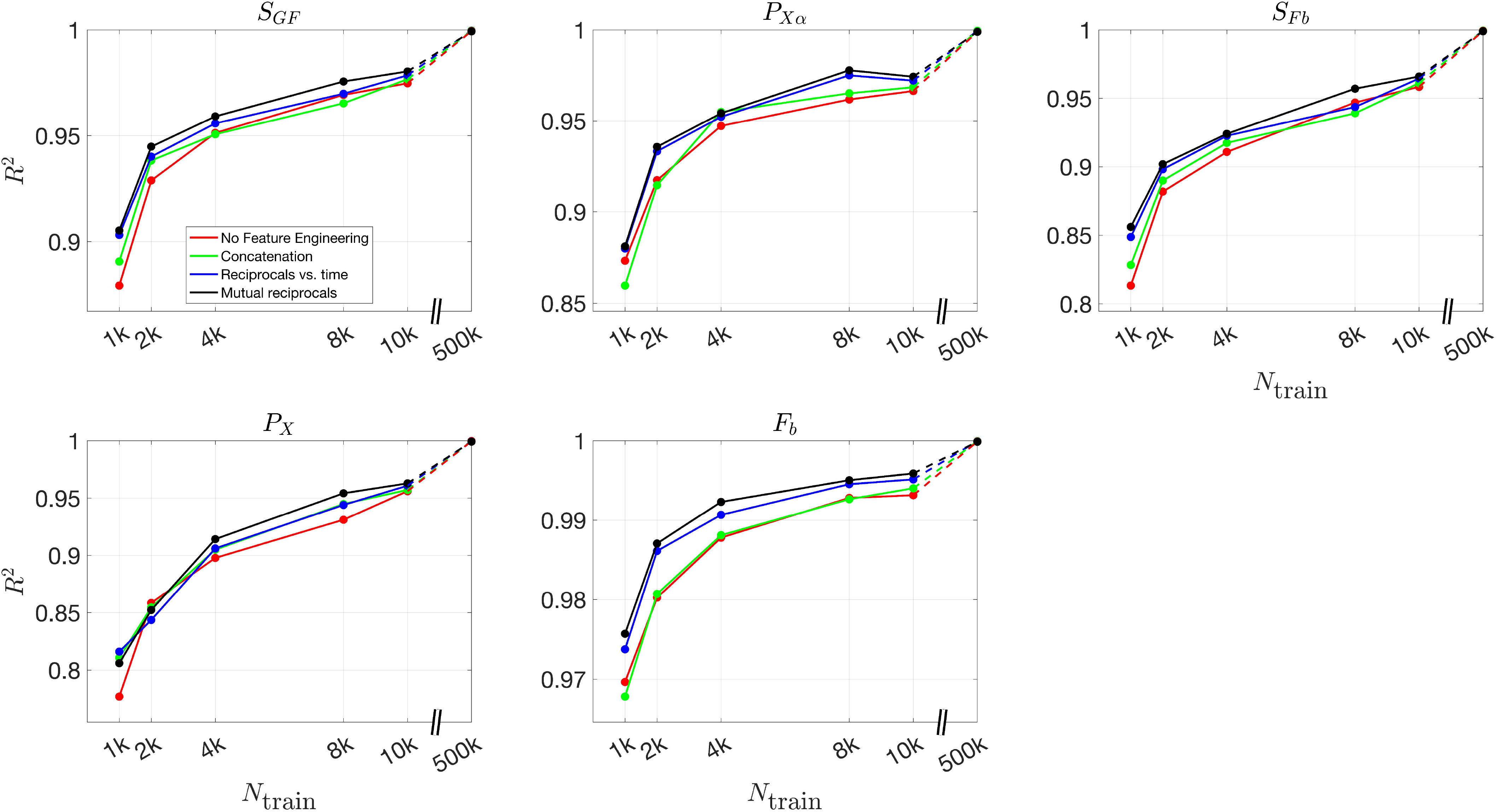

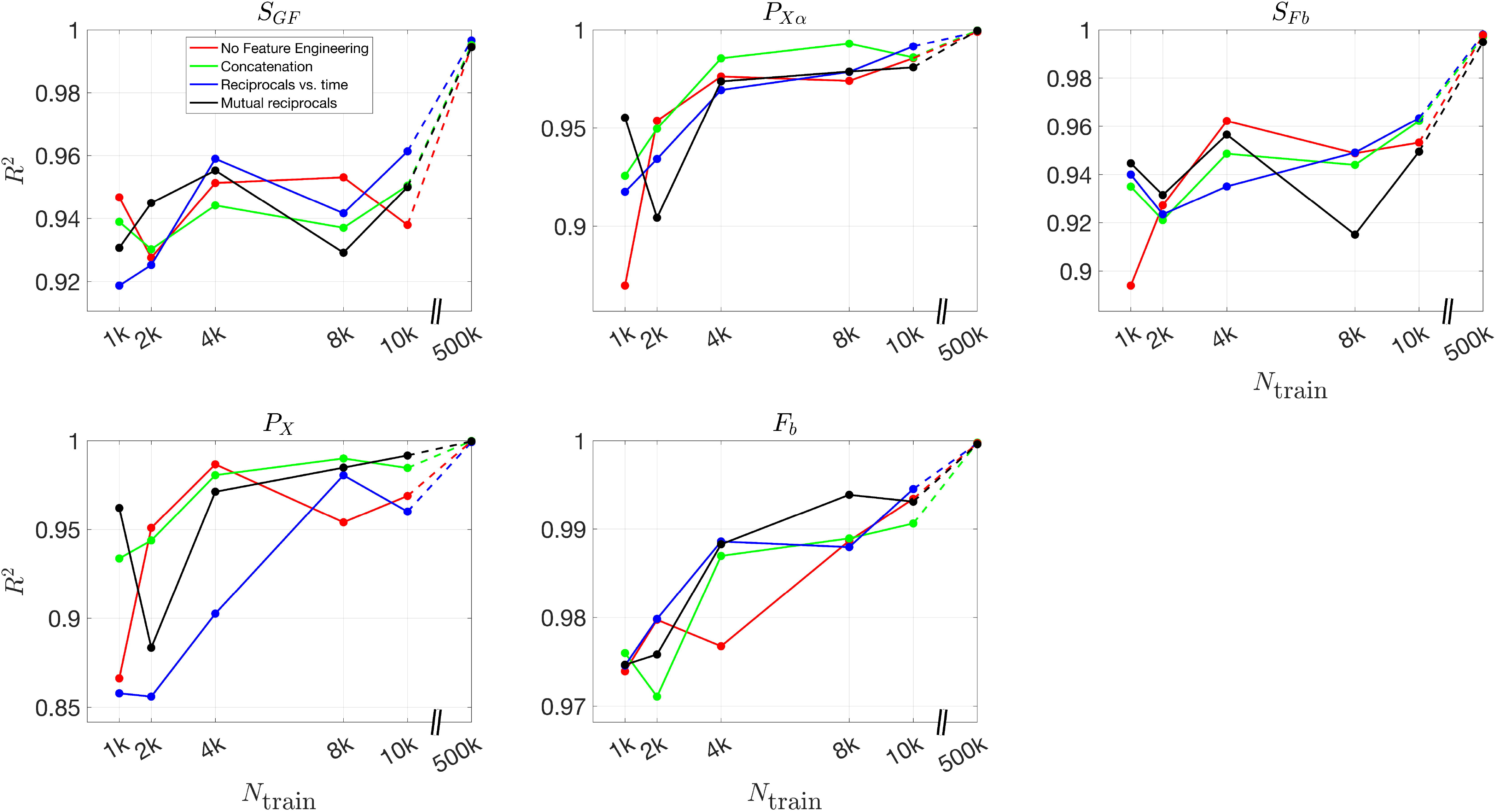

## References

Hornik, Kurt and Stinchcombe, Maxwell and White, Halbert. Multilayer feedforward networks are universal approximators. Neural networks, 1989, 2(5):359–366.

Bergman, Richard N and Ider, Y Ziya and Bowden Charles R and Cobelli, Claudio Quantitative estimation of insulin sensitivity. American Journal of Physiology-Endocrinology And Metabolism

Periwal, Vipul and Chow Carson C and Bergman Richard N and Ricks, Madia and Vega Gloria L and Sumner Anne E. Evaluation of quantitative models of the effect of insulin on lipolysis and glucose disposal American Journal of Physiology-Regulatory, Integrative and Comparative Physiology.

Stefanovski, Darko and Punjabi Naresh M and Boston Raymond C and Watanabe Richard M. Insulin Action, Glucose homeostasis and free fatty acid metabolism: insights from a novel model. Frontiers in Endocrinology. 2021

Boden, Guenther and Shulman, GI. Free fatty acids in obesity and type 2 diabetes: defining their role in the development of insulin resistance and β-cell dysfunction. European journal of clinical investigation. 2002

Savage, David B and Petersen Kitt F and Shulman Gerald I. Mechanisms of insulin resistance in humans and possible links with inflammation. Hypertension. 2005

