## Supplementary Text S1 for "Deep learning approach to parameter optimization for physiological models"

### Text S1: data simulation

#### 1 Outliers of the subject data

Throughout this study, we consider subjects 3,11,18 and 25 as outliers and exclude their physiological data from the procedures described in Section 2. These subjects are excluded due to the distinct characteristics of their FFA data, particularly in terms of monotonicity and range. Including these outliers would complicate the generation of simulated data, the training of the neural network, and the accurate inference of parameters from the trained network. Figure S1 illustrates the FFA physiological data for all 25 subjects. The black curves correspond to subjects 11, 18, and 25, whose FFA values are notably low near the end of the time course. Meanwhile, the brown curve represents subject 3, whose FFA values are disproportionately high over the interval $[50, 80]$.

#### 2 Parameter optimization for models

For the 3D model, the optimization is carried out by minimizing the dimensionless cost function defined as:

$$C_{\mathrm{3D}} = \underset{t \in t_{\mathrm{post20}}}{\mathrm{mean}} \left[ \frac{(G(t) - G_{\mathrm{data}}(t))^2}{G_{\mathrm{data}}(t)^2} + \frac{(F(t) - F_{\mathrm{data}}(t))^2}{F_{\mathrm{data}}(t)^2} \right], \tag{1}$$

where $G(t)$ and $F(t)$ represent the glucose and FFA trajectories, respectively, obtained from the integration of the ODE model using parameter values from each optimization step. $G_{\mathrm{data}}(t)$ and $F_{\mathrm{data}}(t)$ denote the physiological glucose and FFA data. At every iteration, given the $C_X$ value and interpolation of $I_{\mathrm{data}}(t)$, we integrate the equation

$$\frac{\mathrm{d}X}{\mathrm{d}t} = C_X(\max\{I(t) - i_b, 0\} - X(t)), X(0) = 0, \tag{2}$$

up to $t = 22$ to compute $X_{22}$ for the subsequent iteration.

Similarly, for the 2D model, we minimize the cost function

$$C_{\mathrm{2D}} = \underset{t \in t_{\mathrm{all}}}{\mathrm{mean}} \frac{(F(t) - F_{\mathrm{data}}(t))^2}{F_{\mathrm{data}}(t)^2}. \tag{3}$$

Figures S2, S3 and S4 show the fittings of glucose and FFA for all 21 subjects with the 3D and 2D models, respectively.

#### 3 Data simulation with Gaussian Process Regression

The Gaussian Process Regression(GPR) model we use has Radial Quasi-Periodic (RQ) kernel:

$$k_{\mathrm{RQ}}(\mathbf{x}_1, \mathbf{x}_2) = \left(1 + \frac{1}{2\alpha}(\mathbf{x}_1 - \mathbf{x}_2)^{\mathrm{T}} \Theta^{-2} (\mathbf{x}_1 - \mathbf{x}_2)\right)^{-\alpha}, \tag{4}$$

where $\Theta$ is a lengthscale parameter with a prior in $(0.01, 5)$, and the rational quadratic relative weighting parameter has prior $(0.8, 1)$. With this kernel, we define the exact marginal log likelihood as

$$\mathrm{mll}(\mathbf{x}_1, \mathbf{x}_2) = -\frac{1}{2}(\mathbf{x}_1 - \mu)^{\mathrm{T}} K^{-1} (\mathbf{x}_2 - \mu) - \frac{1}{2}\log\det(K) - \frac{n}{2}\log(2\pi), \tag{5}$$

where $K_{ij} = k_{\mathrm{RQ}}(x_1^{(i)}, x_2^{(j)})$. Suppose we have physiological data of insulin from $N$ subjects, normalize them using the mean and standard deviation at each individual time point, and denote the normalized data as $\{I_{\mathrm{Nmlz}}^{(i)}(t) \mid t \in t_{\mathrm{vec}}\}_{i=1}^{N}$, where $t_{\mathrm{vec}} = t_{\mathrm{all}}$ or $t_{\mathrm{post20}}$. During the training of this GPR model $M$, each time we randomly choose a subject index $i$, and calculate the value and backpropagate gradients of the following loss function $L$

$$L = -\mathrm{mll}(M(t_{\mathrm{vec}}), I_{\mathrm{Nmlz}}^{(i)}(t_{\mathrm{vec}})). \tag{6}$$

To maximize the marginal log likelihood, we iterate this process and update the model parameters to minimize $L$ until it is lower than a predefined threshold (we set it as 0.73 for the FSIGT physiological data used in this study). To achieve this, we employ the Adam optimizer with a learning rate of 0.001.

### 4 Sampling of parameter set

Here we put the details of generating sample parameter set. For each parameter $p$, its optimized values are obtained from the data of all subjects. Additionally, before integrating the model, we must generate samples for $G_b$, $F_b$, $G(22)$(initial condition of $G$), and $F(22)$(initial condition of $F$). These values are directly obtained from the physiological data of each subject. In the remaining sections of this paper, we treat $G_b$ and $F_b$ as parameters of 3D model, and $F_b$ as a parameter of the 2D model. Notably, $I_b$ is the first value of the insulin data generated using Gaussian process regression (GPR), so $I_b$ does not require sampling at this stage.

We use a multivariate log-normal distribution to fit the parameter sets from optimization, and generate as many parameter samples as required. Figure S5 and S6 show the correlation between the optimized parameters and generated samples for these two models respectively.

After generating the simulated data, we filter out parameter-trajectory pairs where the glucose or FFA trajectories contain negative values. For the 3D model, we also remove cases where the trajectory is not monotonically decreasing over the interval [22,30], which includes five time points. This filtering step ensures that the dataset remains consistent with expected physiological patterns.
