## Supplementary Text S2 for "Deep learning approach to parameter optimization for physiological models"

### Text S2: Details of the primary neural network

#### 1 Training of the neural network

We generate a simulated dataset using the method described in Section 4 of the main text. The dataset is randomly split, with 80% allocated for training and 20% for validation. To ensure consistency and reproducibility, a fixed random seed is used during the splitting process. The training data is shuffled to prevent order bias, while the validation set remains fixed for consistent evaluation. Mini-batches are employed to enhance computational efficiency and training stability.

A separate testing dataset is generated using the same procedure as the training and validation datasets. This testing dataset is never used during training or validation, ensuring an unbiased assessment of the trained model. The same testing dataset is applied across all models for fair comparison.

Before training, each column of the neural network output $d_O$ is linearly rescaled to the range $[0, 1]$. During the inference step after the training is complete, the rescaling is inversely applied to the outputs to recover the original parameter values.

The neural network is trained using the Adam optimizer with the learning rate Lr schedule given by the following function:

$$\mathrm{Lr(epoch)} = \begin{cases} \mathrm{MaxLr} \times \mathrm{C_3} / \left(1 + \exp\left(-\mathrm{C_1} \cdot (\mathrm{epoch} - \mathrm{N_{top}}/2)\right)\right) & ,\mathrm{epoch} \leq \mathrm{N_{top}}, \\ \mathrm{MaxLr} \times \mathrm{C_4} / \left(1 + \exp\left(-\mathrm{C_2} \cdot \left(\frac{3}{4}\mathrm{N_{max}} - \mathrm{epoch}\right)\right)\right) & ,\mathrm{epoch} > \mathrm{N_{top}}, \end{cases} \tag{1}$$

where the maximal learning rate MaxLr, epoch number $N_{\mathrm{top}}$ where learning rate arrives at maximum, and number of epoch $N_{\max}$ for the training are to be determined, and $C_1 = 10/N_{\mathrm{top}}$, $C_2 = 10/N_{\max}$. $C_3 = 1 + \exp(-C_1 N_{\mathrm{top}}/2)$ and $C_4 = 1 + \exp\left(-C_2\left(\frac{3}{4}N_{\max} - N_{\mathrm{top}}\right)\right)$ are such that this function is continuous at $N_{\mathrm{top}}$. The training process span 1000 epoches, and the model with the lowest validation loss is saved. For the training of the network, we set maximum learning rate of $10^{-3}$, batch size 500, $N_{\max} = 2000$ and $N_{\mathrm{top}} = 1000$.

#### 2 Feature Engineering and neural network architecture

Let $N$ be the batch size. We first denote some layers as follows which we will use in feature engineering:

1. MaxPooling2D, a pooling layer with kernel size=$(1, 2)$;
2. Flatten, input size=$(N, 1, N_1, N_{\mathrm{conv}})$, output size=$(N, N_1 N_{\mathrm{conv}})$;
3. Dense$_1$: dense layer with input size same as the output size of Flatten layer, and output size=$(N, N_2)$, employing the hyperbolic tangent function (tanh) as its activation function;
4. Dense$_2$: dense layer with input size $(N, N_2)$ and output size=$(N, N_3)$, employing the hyperbolic tangent function (tanh) as its activation function;
5. Dense$_3$: the final dense layer to output $d_O$, with input size=$(N, N_3)$ and output size=$(N, N_{\mathrm{para}})$. It uses ReLU or $\frac{1}{2}(\tanh(x) + 1)$ as final activation function, considering that all parameters are positive before and after rescaling.

Here we show the details of several features:

1. No Feature Engineering:

   $d_I$ is processed by a two-dimensional convolutional layer (conv2D) called conv2D$_1$ with kernel size=(2,2) and stride=(2,1), followed by a Maxpooling2D layer. Then it is processed by another 2D convolutional layer called conv2D$_2$ with kernel size=(6,2), followed by a Maxpooling2D layer. The data is then flattened and fed into two dense layers Dense_1 and Dense_2. The input and output sizes of these layers are put in the Table 1.

| 2D model | conv2D$_1$ | conv2D$_2$ | Dense$_1$ | Dense$_2$ |
|---|---|---|---|---|
| input size | $(N, 12, 28, 1)$ | $(N, 6, 13, 1)$ | $(N, 3072)$ | $(N, 1024)$ |
| output size | $(N, 6, 27, 1)$ | $(N, 1, 12, 1)$ | $(N, 1024)$ | $(N, 1024)$ |
| 3D model | conv2D$_1$ | conv2D$_2$ | Dense$_1$ | Dense$_2$ |
| input size | $(N, 12, 16, 1)$ | $(N, 6, 7, 512)$ | $(N, 1536)$ | $(N, 1024)$ |
| output size | $(N, 6, 15, 512)$ | $(N, 1, 6, 512)$ | $(N, 1024)$ | $(N, 1024)$ |

**Table 1.** Input and output sizes for layers for the 2D and the 3D models for feature 1: no feature engineering. The four numbers for conv2D layers represent, respectively, the batch size, the numbers of rows, the numbers of columns, and the layer dimension.

2. Concatenation: $d_I$ has the same structure as in Feature 1 (No Feature Engineering), with an additional concatenation process: the input data $d_I$ is initially processed by a two-dimensional convolutional layer conv2D$_1$, with kernel size=(2,2), stride=(2,1), using padding of replication on the right. Subsequently, its output is concatenated with $d_I$ to incorporate the functions of time and substances (glucose, FFA, and insulin), as well as their derivatives with respect to time. This concatenation is performed by pairing the 12 rows of $d_I$ into 6 pairs, and after each pair, inserting the corresponding row from the output of conv2D$_1$. The concatenated data then passes through two conv2D layers (denoted as conv2D$_2$ and conv2D$_3$, respectively) each followed by a Maxpooling2D layer. conv2D$_2$ has kernel size=(3,2), stride=(3,1); conv2D$_3$ has kernel size=(6,2). The data is then flattened and fed into two dense layers as before. The full process is illustrated in Figure S9, and the input and output sizes of layers are put in Table 2.

| 2D model | conv2D$_1$ | conv2D$_2$ | conv2D$_3$ | Dense$_1$ | Dense$_2$ |
|---|---|---|---|---|---|
| input size | $(N, 12, 28, 1)$ | $(N, 18, 28, 512)$ | $(N, 6, 27, 512)$ | $(N, 3072)$ | $(N, 1024)$ |
| output size | $(N, 6, 28, 512)$ | $(N, 6, 27, 512)$ | $(N, 1, 13, 512)$ | $(N, 1024)$ | $(N, 1024)$ |
| 3D model | conv2D$_1$ | conv2D$_2$ | conv2D$_3$ | Dense$_1$ | Dense$_2$ |
| input size | $(N, 12, 16, 1)$ | $(N, 18, 16, 512)$ | $(N, 6, 15, 512)$ | $(N, 1536)$ | $(N, 1024)$ |
| output size | $(N, 6, 16, 512)$ | $(N, 6, 15, 512)$ | $(N, 1, 7, 512)$ | $(N, 1024)$ | $(N, 1024)$ |

**Table 2.** Input and output sizes for layers for the 2D and the 3D models for feature 2: concatenation. The four numbers for conv2D layers represent, respectively, the batch size, the numbers of rows, the numbers of columns, and the layer dimension.

### 3 Denoising from physiological data

The neural network we train takes trajectory data of ODE systems as input. However, real physiological data are not smooth like the model-generated solutions. Therefore, it is necessary to denoise the physiological data before using it as input for the neural network. To achieve this, we employ a simple fully connected neural network for denoising, and prepare the training data as follows:

- The output of the network is the simulated trajectory set obtained in the earlier section.
- The input for each output trajectory is generated by adding noise to the simulated trajectory data.

Here we show how we extract adjacent noise information from the physiological data, by taking the FFA data over $N$ time points as an example. For subject $k$, let

$$\Delta F(k,l) = \begin{cases} \left| \frac{F_{\text{data}}(k,l+1)-F_{\text{data}}(k,l)}{F_{\text{data}}(k,l+1)+F_{\text{data}}(k,l)} \right|, & \text{if } l = 1, \\ \frac{1}{2}\left[ \left| \frac{F_{\text{data}}(k,l+1)-F_{\text{data}}(k,l)}{F_{\text{data}}(k,l+1)+F_{\text{data}}(k,l)} \right| + \left| \frac{F_{\text{data}}(k,l)-F_{\text{data}}(k,l-1)}{F_{\text{data}}(k,l)+F_{\text{data}}(k,l-1)} \right| \right], & \text{if } 2 \le l \le N-1, \\ \left| \frac{F_{\text{data}}(k,l)-F_{\text{data}}(k,l-1)}{F_{\text{data}}(k,l)+F_{\text{data}}(k,l-1)} \right|, & \text{if } l = N \end{cases} \tag{2}$$

denote the adjacent increment at the $l$-th point over the time course. We add 20 percent noise to each of the output of the neural network, using the following formula:

$$F_{\text{noise}}(k,l) = \begin{cases} F_{\text{data}}(k,l), & \text{if } l = 1, \\ F_{\text{data}}(k,l) + 0.2Y\left[F_{\text{data}}(k,l-1) + F_{\text{data}}(k,l)\right], & \text{if } l \ge 2, \end{cases} \tag{3}$$

where $Y \sim \mathcal{N}(0, \underset{k}{\text{mean}}\,(\Delta F(k,l)))$ is sampled from a normal distribution with a mean of 0 and a standard deviation equal to the mean of adjacent increment $\Delta F(k,l)$ across all subjects $k$. This noisy trajectory serves as the corresponding input data for training the neural network.

We use the FFA data from the first 200,000 training samples as the output and add noise to these samples using the above method to generate the corresponding input data. A fully connected neural network is then trained with this data. The network consists of three dense layers, each with 256 nodes followed by the ReLU activation function. The loss function is set to be the mean squared error (MSE) between the inferred outputs and the given outputs. The network is trained using the learning rate schedule defined by (1), with $N_{\text{max}} = 100$ and $N_{\text{top}} = 50$.

Once the training is complete, we feed the glucose and FFA physiological data from all subjects into the trained network, which outputs the corresponding denoised data. Figure S8 illustrates the denoised FFA data for all subjects.
